# Host Species is Linked to Pathogen Genotype for the Amphibian Chytrid Fungus (*Batrachochytrium dendrobatidis*)

**DOI:** 10.1101/2021.11.24.469827

**Authors:** Allison Q. Byrne, Anthony W. Waddle, Veronica Saenz, Michel Ohmer, Jef R. Jaeger, Corinne L. Richards-Zawacki, Jamie Voyles, Erica Bree Rosenblum

## Abstract

Host-pathogen specificity can arise from certain selective environments mediated by both the host and pathogen. Therefore, understanding the degree to which host species identity is correlated with pathogen genotype can help reveal historical host-pathogen dynamics. One animal disease of particular concern is chytridiomycosis, typically caused by the global panzootic lineage of the amphibian chytrid fungus (*Batrachochytrium dendrobatidis*, Bd), termed the Bd-GPL. This pathogen lineage has caused devastating declines in amphibian communities around the world. However, the origin of Bd-GPL and the fine-scale transmission dynamics of this lineage have remained a mystery. This is especially the case in North America where Bd-GPL is widespread, but disease outbreaks occur sporadically. Herein, we use Bd genetic data collected throughout the United States from amphibian skin swab and cultured isolate samples to investigate Bd genetic patterns. We highlight two case studies in Pennsylvania and Nevada where Bd-GPL genotypes are strongly correlated with host species identity. Specifically, in some localities bullfrogs (*Rana catesbeiana*) are infected with Bd-GPL lineages that are distinct from those infecting other sympatric amphibian species. Overall, we reveal a previously unknown association of Bd genotype with host species and identify the eastern United States as a *Bd* diversity hotspot and potential ancestral population for Bd-GPL.

## Introduction

When host-pathogen specificity exists within a system it can serve to reveal historical disease dynamics and has consequences for the evolution of virulence. Theoretical work predicts host-pathogen specificity will evolve, as a pathogen consistently exposed to a particular host would be subject to stronger selection from that host and therefore evolve faster than a generalist pathogen (1). However, the high frequency of multi-host pathogens in nature implies that interacting factors present in complex natural systems may tip the scales in favor of generalist pathogens (2). When host-specificity is high, it is predicted that host-pathogen feedback is predicted to favor selection towards decreased pathogen virulence because an increase in virulence would be disadvantageous for the preferred host (3). Similarly, low levels of specificity favor pathogens with high virulence and often lead to the competitive exclusion of other less fit pathogen strains (3). Therefore, understanding the degree to which host-pathogen specificity exists in a system can shed light on the underlying evolutionary processes and disease dynamics at play.

Advances in genetic monitoring tools have allowed us to investigate fine scale disease dynamics in natural systems and uncover previously unknown host-pathogen genetic associations. The utility of DNA sequence data in understanding pathogen evolution and genetic diversity has been evidenced by genetic monitoring efforts during the Covid-19 pandemic (e.g., (4). By applying technologies used in human pathogen systems to wildlife diseases we can investigate the epidemiology of diseases that are of conservation concern. Of particular interest is the amphibian chytrid fungus [*Batrachochytrium dendrobatidis* (Bd)] (5) because of its association with amphibian die offs and extinctions in many disparate areas of the world (6–10). For Bd, there are at least five distinct genetic lineages, but the Global Panzootic Lineage (Bd-GPL) has been associated with almost all disease outbreaks around the world (11–13). Recent DNA sequencing of Bd samples collected in North America has confirmed previous reports of two distinct sub-lineages within the Bd-GPL, termed “Bd-GPL1” and “Bd-GPL2” (14–16). While it has become increasingly clear that there is deep genetic structure within the Bd-GPL, and that both Bd-GPL1 and Bd-GPL2 often co-occur, what remains unclear is how these patterns arose and are maintained across fine spatial scales.

A growing number of studies have shown that Bd, and specifically the Bd-GPL, has patterns of genetic diversity that are linked to geography, but none have tested the hypothesis that there is a link between Bd genetics and host species remains untested (13,15,16). In this study, we investigated the relationship between host species and Bd genotype in two distinct amphibian communities within the United States. One community was sampled in northwestern Pennsylvania near the Pymatuning Lab of Ecology (PA). Sampling in this temperate region took place at eight sites (including permanent and ephemeral ponds and one forested stream site). The second amphibian community was sampled in southern Nevada (NV) within the eastern Mojave Desert in an area characterized by permanent water bodies separated by larger distances. While most amphibian species encountered in this study were found in only one of these regions, the American bullfrog [*Rana catesbeiana* (Shaw 1802), hereafter “bullfrogs”] occurred across both regions in either its native (PA) or introduced (NV) range. By comparing the relationship between Bd genotype and host species in these two localities we not only reveal more about the evolutionary history of Bd in the United States (US), but can also begin to understand the repeatability of these patterns across systems and the role certain amphibian species may have in shaping Bd genetic diversity.

## Materials and Methods

### Sample Collection in Northwestern Pennsylvania and Southern Nevada

We collected Bd samples in PA between 2017–2019 and in NV between 2016–2018. In both areas, we captured frogs by hand using clean gloves. In PA we also used nets with recommended field hygiene protocols followed (17). We kept captured amphibians individually in clean plastic bags during processing. To collect skin cell samples, we swabbed frogs in PA 5 times each on dorsal, ventral, right, and left sides, and feet with a sterile swab. In NV, we swabbed frogs 10 times on each ventral side and 5 times on each foot (18). We initially stored all swabs in tubes on ice and promptly transferred them to a freezer at -20 °C until processing. We extracted DNA from swabs in PA using the Qiagen DNEasy Blood and Tissue kit (Qiagen,Valencia, USA), and from NV using PrepMan Ultra (19), following both manufacturer’s protocols.

To generate pure cultures of Bd from the relict leopard frog (*Rana onca*) we used a non-lethal method in the field (20). We excised a small piece of webbing tissue (1 mm^2^) that was cleaned and embedded into TGhL agar plates containing a cocktail of antibiotics (kanamycin, ciprofloxacin, streptomycin, and penicillin) (21). For the other species in NV from which cultures were derived, we brought the animals into the laboratory and humanely euthanized them prior to excising 1 mm^2^ pieces of skin tissue from the ventral abdomen, legs, and rear feet. Each tissue sample was then wiped through and embedded into a TGhL agar plate containing antibiotics. We established pure cultures by transferring Bd growth to H-broth liquid medium. We used a modified phenol-chloroform extraction to isolate DNA from the Bd cultures (22).

### DNA Sequencing and Preprocessing

We genotyped Bd from PA and NV using a custom genotyping assay (23). For the PA samples, we initially sequenced 115 skin swabs and for the NV samples, we initially sequenced 28 pure isolates and 36 skin swabs. After downstream filtering for missing data (see below) our final dataset included 71 samples from PA, and 52 samples from NV. For some of our analyses we also included previously published Bd sequence data from samples collected from around the world (13).

We used the Fluidigm Access Array platform to perform microfluidic multiplex PCR on 191 regions of the Bd genome and one diagnostic locus for the closely related *Batrachochytrium salamandrivorans* (24). Each target locus is 150-200 base pairs long, with the Bd targets distributed across the Bd nuclear and mitochondrial genomes (23). Extracted DNA from swab samples was cleaned and concentrated using an isopropanol precipitation. Isolate samples were diluted to a concentration of 3ng/µl using PCR-grade water. All samples were preamplified in two separate PCR reactions, each containing 96 primer pairs at a final concentration of 500nM. For each preamplification PCR reaction we used the FastStart High Fidelity PCR System (Roche) at the following concentrations: 1x FastStart High Fidelity Reaction Buffer with MgCl_2_, 4.5mM MgCl_2_, 5% DMSO, 200µM PCR Grade Nucleotide Mix, 0.1 U/µl FastStart High Fidelity Enzyme Blend. We added 1µl of cleaned DNA to each preamplification reaction and used the following thermocycling profile: 95°C for 10 min, 15 cycles of 95°C for 15 sec and 60°C for 4 min. Preamplified products were treated with 4µl of 1:2 diluted ExoSAP-it (Affymetrix Inc.) and incubated for 30 min at 30°C, then 15 min at 80°C. Treated products were diluted 1:5 in PCR-grade water. The diluted products from each of the two preamplification reactions were combined in equal proportions and used for downstream amplification and sequencing. Each preamplified sample was then loaded onto a Juno™ LP 192.24 IFC (Fluidigm, Inc.) for library preparation. This flow cell performs microfluidic multiplex PCR to amplify all samples using 24 separate pools each containing 8 primer pairs. Barcoded samples were then pooled and sequenced on an Illumina MiSeq lane using the 300bp paired-end kits at the University of Idaho IBEST Genomics Resources Core.

### Sequence Data Analysis

We pre-processed all sequencing data as described in (23) and generated consensus sequences for all variants present for each sample at each locus using IUPAC ambiguity codes for multiple alleles. We filtered reads by selecting sequence variants that were present in at least 5 reads and represented at least 5% of the total number of reads for that sample/locus using dbcamplicons (https://github.com/msettles/dbcAmplicons).

We used a gene-tree to species-tree approach to construct phylogenies separately for the PA and NV sequence datasets. These trees allow us to explore the relationship of the Bd collected for this study to previously-published Bd samples representing all known Bd lineages (N=13; 13,25). First, we trimmed our consensus sequence dataset to eliminate loci that had more than 50% missing data for either the NV or PA datasets, resulting in 174 loci. Next, for each dataset we individually aligned all loci using the MUSCLE package (26) in R (v.3.4.3), checked the alignments for errors in Geneious (v.10.2.3) (27), and used the RAxML plugin (28) in Geneious to search for the best scoring maximum likelihood tree for each locus using rapid bootstrapping (100 replicates). We then collapsed all branches in all trees with <10 bootstrap support and used Astral-III to generate a consensus tree (29). Astral generates an unrooted species tree given a set of unrooted gene trees.

To further explore the genetic variation of Bd collected at our sites, we generated a dataset of single nucleotide polymorphisms (SNPs). First, we aligned all raw sample reads (PA and NV) and reads from 26 previously published Bd-GPL reference samples (13,25) using bwa mem (30) to a reference fasta containing target sequences for each amplicon extracted from reference genome JEL423 (Broad Institute). We then used Freebayes (v.1.1.0) to call variants based on haplotypes (31) stipulating that variants only be called on the 174 amplicons that passed the 50% missing data cutoff described above. Variants were then filtered using vcftools (v.4.2) (32) to only include variants with a minor allele frequency > 0.01, quality > 30, less than 10% missing data, and minimum depth of 5. The final variant set included 129 binary SNPs with 3.8% missing data across all samples.

We then imported our SNPs into R using the vcfR package (v.1.12.0) (33) and converted the variants to a genlight object. To determine the optimal number of clusters in our data we used a Discriminant Analysis of Principal Components (DAPC) (34). First, we ran the find.clusters function from the R package adegenet (v.2.1.1) with a maximum number of clusters = 10 (35). This function implemented the clustering procedure used in DAPC by running successive K-means with clusters from K = 1 to K = 10 after transforming the data using a principal component analysis (PCA). For each model BIC was computed as a measure of statistical goodness of fit. The best K was chosen as the value for K after which increasing K no longer leads to an appreciable improvement of fit (measured as substantial decrease in BIC) (34). We then ran the DAPC analysis specifying the optimal number of clusters. To determine the optimal number of PCs to use for this analysis we ran the adegenet function optim.a.score which calculates the alpha score as (P_t_-P_r_), where P_t_ is probability of reassignment to the true cluster and P_r_ is the probability of reassignment to randomly permuted clusters. We used a minimum of 2 PCs in the DAPC calculations. Finally, we conducted a PCA using the glPCA function in adegenet and plotted the PCA using the DAPC clusters to visually group samples together. Samples with posterior probability of inclusion in one cluster < 0.99 were labeled “unassigned.” We plotted the first two PCs which together explain 68.2% of the variation in our data. We then computed a PCA separately for each locality to show within locality diversity across species and sites.

To evaluate the relative contribution of sampling site, host species, and individual variation, we ran an analysis of molecular variance (AMOVA) (36) using our SNP data. This analysis was conducted separately for the PA and NV datasets and determined the relative influence of within individual variation, host species identity, and individual sampling locality on Bd genetic structure. We ran this analysis using the poppr package in R (37) and only used SNPs with <5% missing data. This resulted in 40 high quality SNPS for PA and 126 high quality SNPs for NV. We then tested for statistical significance using the “randtest” function in the package ade4, randomly permuting our data 999 times.

To evaluate the frequency of mixed infections on individuals, we searched our set of amplicon sequences for those that had diagnostic alleles that discriminate between the Bd-GPL1 and Bd-GPL2 lineages. First, we searched the amplicon sequences from the set of 26 pure culture, reference samples. We identified amplicons that had at least 1 SNP that distinguished GPL1 samples from GPL2 (as defined by both the DAPC and phylogenetic analyses described above). The GPL1 Bd isolates in this reference set were CJB4, JEL238, JEL267, JEL359, MexMkt, MLA1, SRS812, TST75. The GPL-2 Bd isolates were Campana, CLB5-2, CJB7, EV001, JAM81, JEL271, JEL275, JEL310, JEL427, JEL429, JEL433, JEL627, Lb_Aber, NBRC106979, Pc_CN_JLV, and RioMaria. These samples were all sequenced from pure cultures grown in the lab, and previously-published whole genome data indicates these are pure Bd cultures representing a single lineage (25,38). From these samples we identified 24 loci that occur on five separate Bd chromosomes that have specific Bd-GPL1 or Bd-GPL2 alleles. For the diagnostic alleles on chromosome 1, heterozygotes (those having both alleles present) are characteristic of Bd-GPL2 isolates (see Figure S3). We then manually scored each sample from PA and NV at each as having either the Bd-GPL1, Bd-GPL2, or both alleles present (heterozygote).

## Results

We found a repeated pattern of Bd genetic structure that is most closely associated with amphibian host species in both PA and NV. The DAPC analysis of the combined PA and NV Bd genetic data show that two clusters (K=2) was the most likely grouping for our data (as inferred from ΔBIC, Figure S1B). In the combined PCA, we see a clear division of Bd genotypes into two clusters, with the presence of intermediate “unassigned” samples (Figure 1C). By evaluating each site separately, we see that PA harbors much more genetic variability within samples (accounting for all “unassigned” Bd genotypes, Figure 1). We confirmed that “unassigned” samples were not simply the product of missing data by showing that each genotype group (Bd-GPL1, Bd-GPL2, and unassigned) had the same distribution of missing data (Figure S1 A,C).

**Figure 1:**
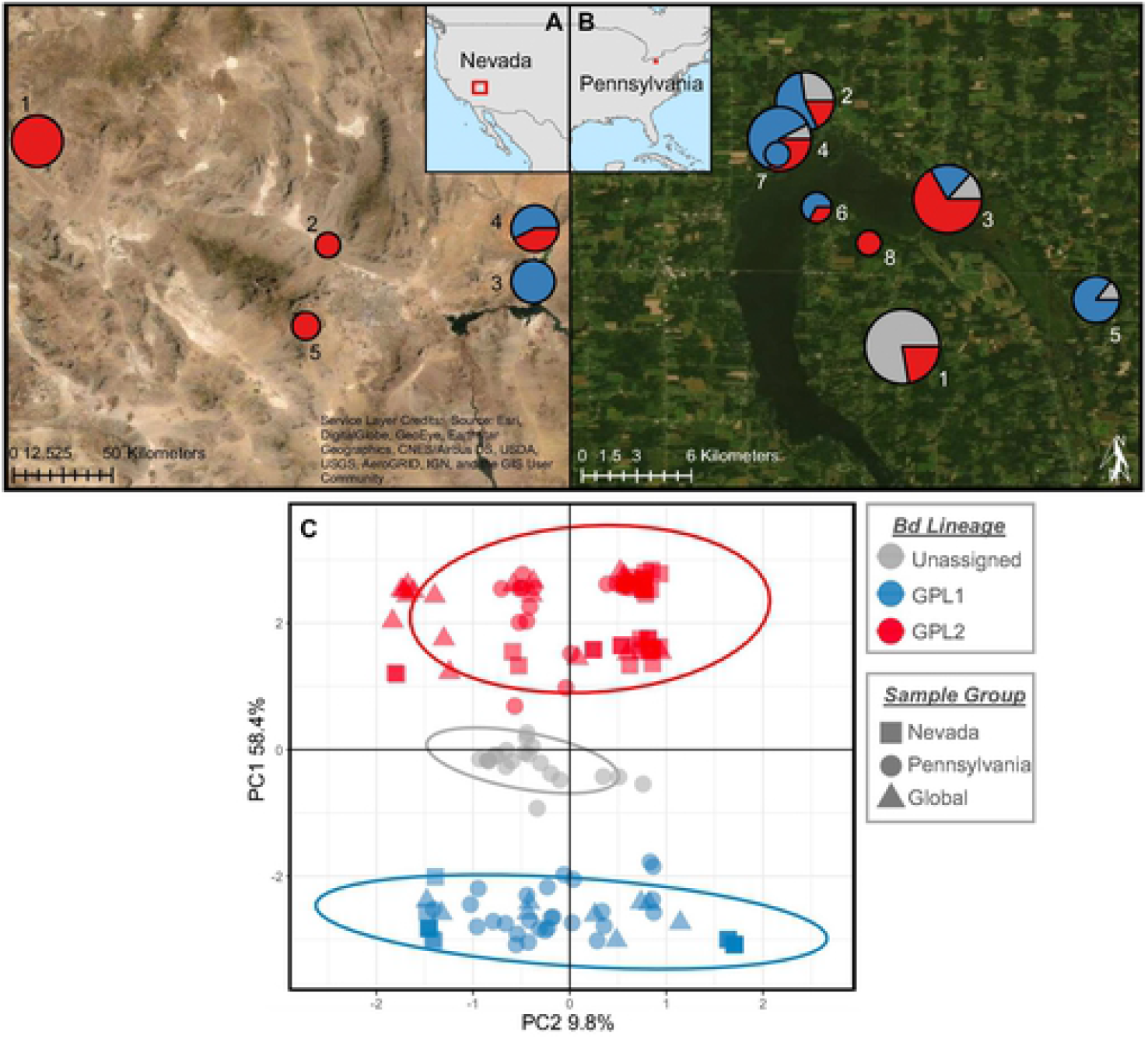
Map of sampling localities for Bd in southern Nevada (A) and northwestern Pennsylvania (B). Individual sampling localities are numbered and represented with a pie chart that shows the proportion of samples belonging to each Bd-GPL sub-lineage as determined using the DAPC clustering analysis. Unassigned samples (in grey) are those with a posterior probability of assignment to either group < 0.99. Pie charts are scaled in size according to the number of samples collected at that site. (C) PCA calculated from 129 SNPS using the combined dataset of southern Nevada (N=52, squares), northwestern Pennsylvania (N=71, circles), and global reference samples (N=26, triangles). Symbols and ellipses colored as in maps above.

Further phylogenetic analyses incorporating previously published Bd isolates shows that that all Bd samples sequenced for this study belong to the Bd-GPL lineage (Figures 2,3) and that the two clusters identified in our dataset correspond to the previously published Bd-GPL subclades Bd-GPL1 and Bd-GPL2 (14,39). The difference between Bd-GPL1 and Bd-GPL2 samples is shown clearly in the combined PCA (Figure 1C), where PC1 (which explains 58.4% of the variation in our genetic data) separates samples into each of these subclades. In NV, 20 samples were assigned to Bd-GPL1 and 32 samples were assigned to Bd-GPL2 (Figure 2). In PA, 27 samples were assigned to Bd-GPL1, 23 samples were assigned to GPL-2, and 21 samples we unassigned (Figure 3).

**Figure 2:**
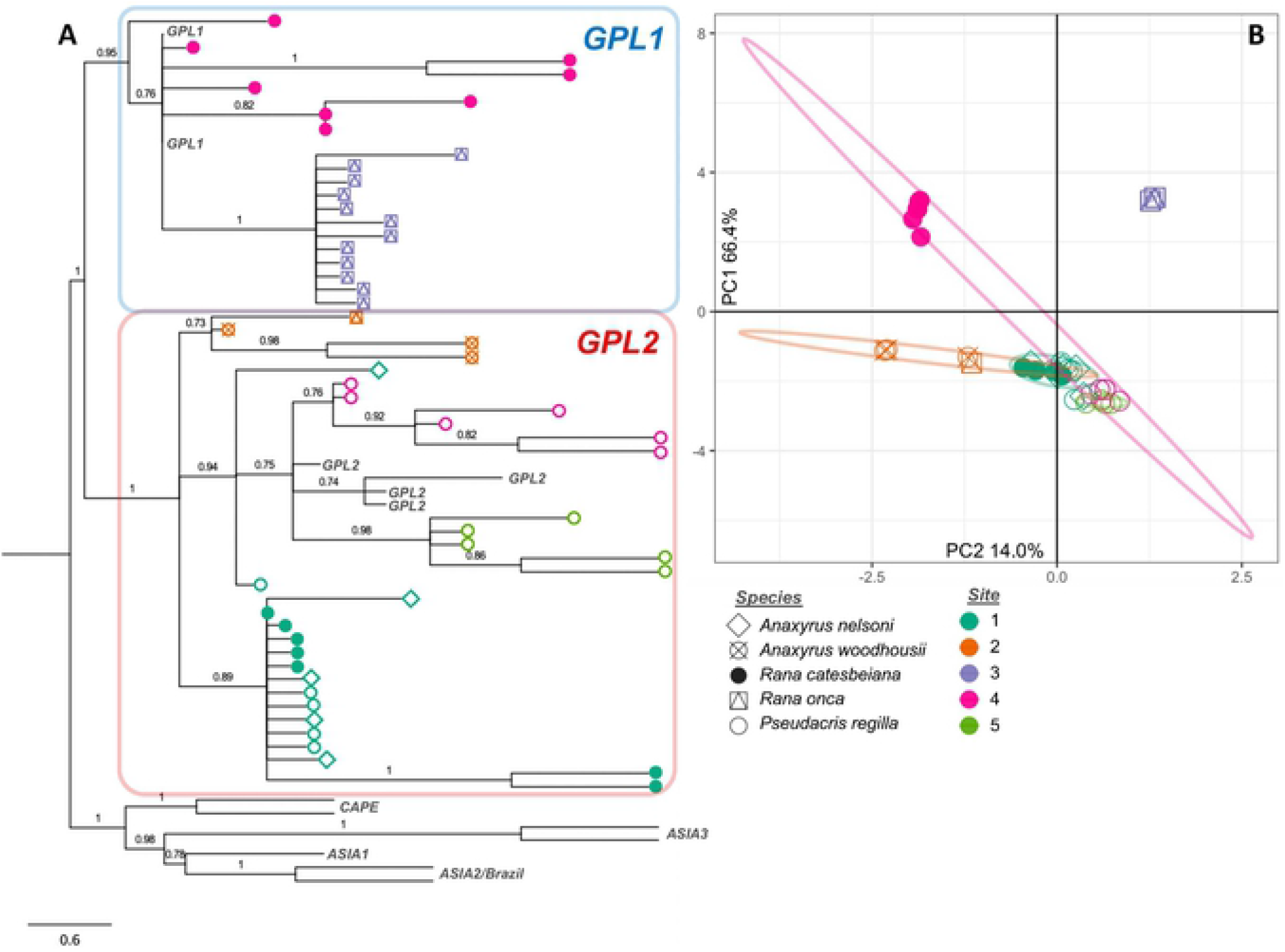
(A) Phylogenetic tree and (B) PCA for 52 Bd samples collected in southern Nevada. Colors represent the locality where the Bd was sampled (numbered as in Figure 2) and symbols represent host species (as in Figure 2). Tree is calculated using Astral-III and includes and additional 13 reference samples representing all known clades of Bd. Nodes are labeled with posterior probability and those with posterior < 0.7 have been collapsed.

**Figure 3:**
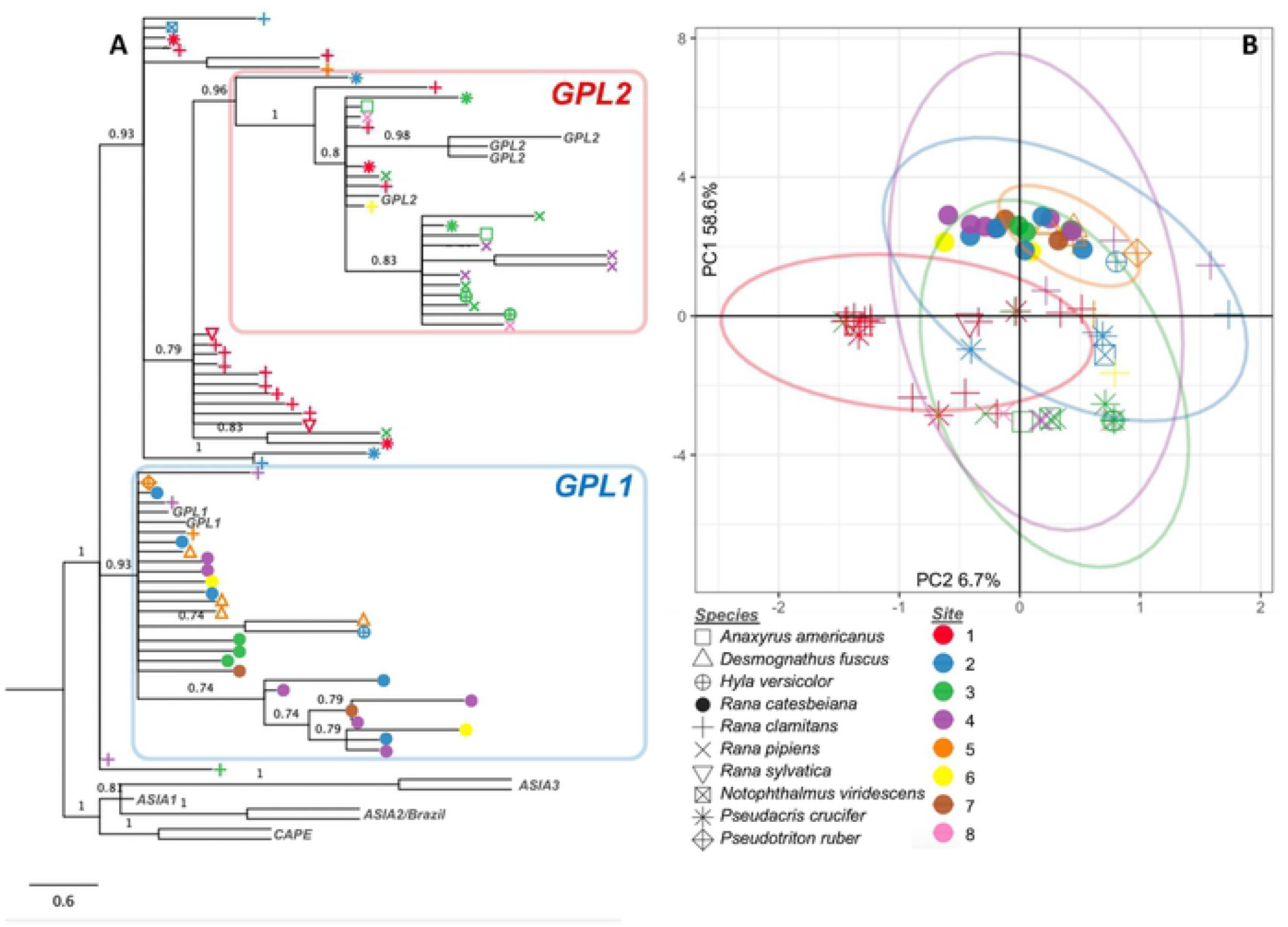
(A) Phylogenetic tree and (B) PCA for 71 Bd samples collected in northwestern Pennsylvania. Colors represent the locality where the Bd was sampled (numbered as in Figure 2) and symbols represent host species (as in Figure 2). Tree is calculated using Astral-III and includes and additional 13 reference samples representing all known clades of Bd. Nodes are labeled with posterior probability and those with posterior < 0.7 have been collapsed.

The factors that may explain Bd genetic structure at each site were determined using an AMOVA. We found that most of the variation in Bd genotypes (64.3% in PA, 58.4% in NV) can be explained by variation between host species, rather than within an individual (34.7% in PA, 8.8% in NV) or between sampling sites (1.0% in PA, 32.8% in NV). Comparisons of variation within individuals and variation between host species were statistically significant using permutation tests to randomly permute samples between groups (999 times, p < 0.001 for each test, Figure S2).

We can evaluate each individual sample for signatures of coinfections or the presence of a hybrid lineage (i.e., the presence of both Bd-GPL1 and Bd-GPL2) by looking at Bd-GPL1/2 diagnostic alleles. The allelic patterns show that the “unassigned” samples in PA appear to either be from a mixed infection or infection with a hybrid lineage – that is, they show a high frequency of heterozygosity and/or the presence of alleles from different Bd-GPL subclades at genomically proximate loci. In PA, one site (PA Site 1) accounts for most of the unassigned samples (67%, 14/21); additionally, 71% (15/21) of unassigned Bd samples were collected from green frogs (*Rana clamitans*). Furthermore, all 18 samples collected from bullfrogs in PA were assigned to GPL-1, despite being collected at five different localities, each with multiple Bd-GPL genotypes present (Figure 3, Figure 4).

**Figure 4:**
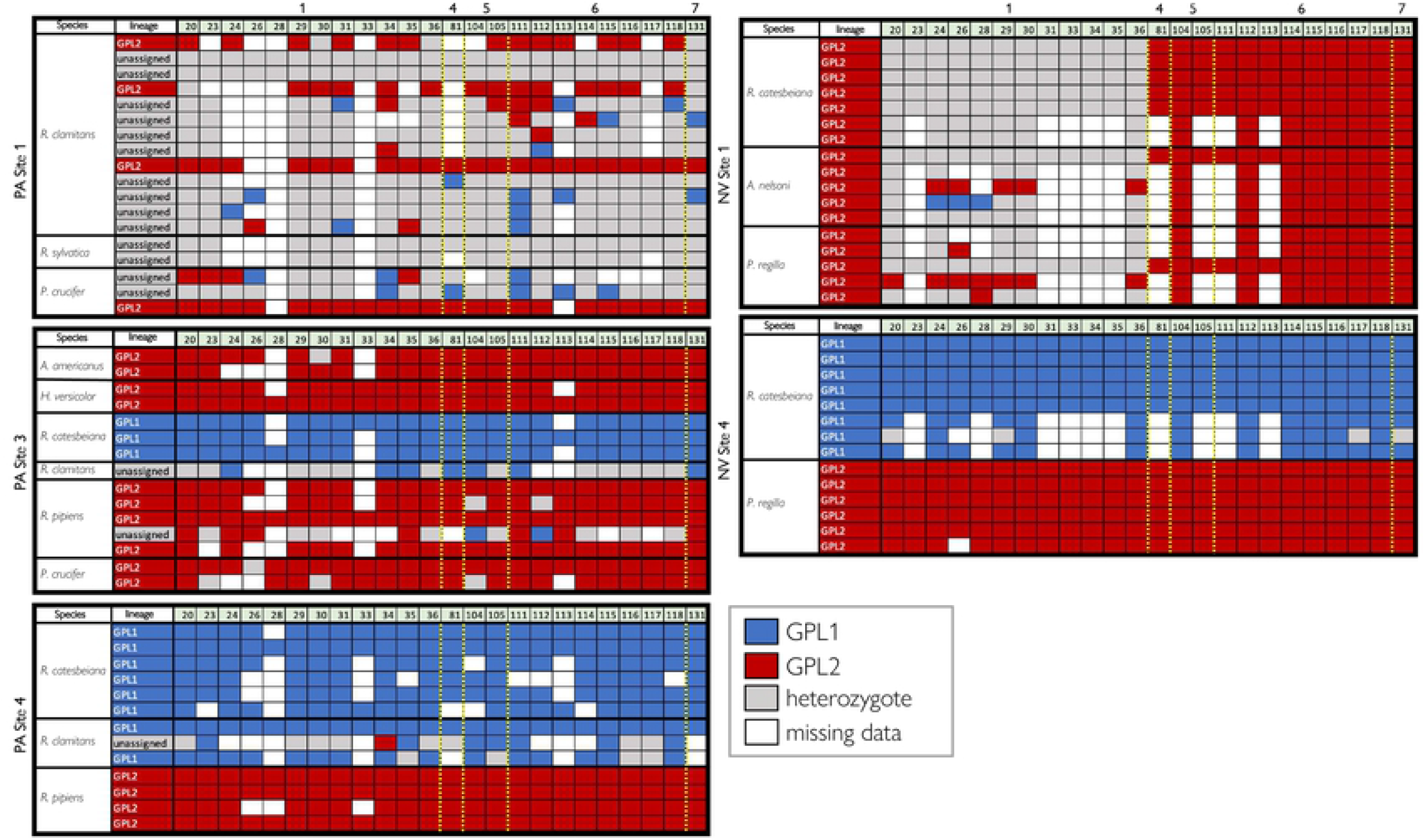
Alleles for PA Site 1 (left top), PA Site 3 (left middle), PA Site 4 (left bottom), NV Site 1 (right top), and NV Site 4 (right bottom). Each row represents an individual sample, and each numbered column represents a nuclear locus that has an informative GPL1/GPL2 allele (N=24). Loci are organized according to position along the Bd genome, with the yellow dashed indicating a change to a new chromosome and the chromosome number labeled along the top. Samples are grouped by host species (“species” column) and labeled with the DAPC assigned lineage (“lineage” column). A red box indicates a Bd-GPL2 allele for that sample at that locus, a blue box represented the Bd-GPL1 allele, a gray box indicates both alleles are present, and a white box indicates missing data. Representative sequences for each allele are included in the supplemental materials.

In NV we see very little Bd-GPL1/Bd-GPL2 mixing within samples and a strong association of the Bd-GPL1 lineage with bullfrogs at one site (NV Site 4), and *R. onca* at another (NV Site 3). While *R. onca* was the only species caught at NV Site 3, bullfrogs were caught alongside Pacific tree frogs (*Pseudacris regilla*) at NV Site 4 and these two species were host to distinct Bd-GPL genotypes with virtually no allele mixing (Figure 4). At another site (NV Site 1), however, bullfrogs were host to Bd-GPL2 as were all other amphibian hosts at that locality. Additionally, the Bd-GPL2 lineage found at NV Site 1 shows a pattern of high heterozygosity at diagnostic alleles found on Bd Chromosome 1, a pattern similar to the reference Bd-GPL2 isolates JEL627, JEL271, Lb_Aber, CJB7, and JAM81 (Figure S3).

## Discussion

Herein, we show a clear, repeated pattern of Bd genetic diversity associated with host species at single geographic sites. While previous studies have shown some broad geographic patterning in Bd genetic structure (15,16), this is the first study to link fine-scale Bd genetic variation to host species identity. Furthermore, this pattern is repeated across individual sites in PA, and in two separate regions of the US. Remarkably, the same association of Bd-GPL1 with bullfrog hosts was seen in both PA and NV. Below, we explore how this pattern may have arisen in each region, discuss what this may mean for the origin of the Bd-GPL, and explore the potential importance of genetically diverse Bd infections.

### How did host-pathogen specificity arise in each region?

Processes occurring at different timescales could produce the patterns of Bd genotype and host species association observed in each region. One possibility is that long-term coevolutionary relationships between Bd and its host species likely triggered *in situ* evolution and subsequent pathogen divergence over time in the ancestral Bd-GPL population. This process could have been mediated by many factors including host immune systems, skin microbial communities, and the environment. Furthermore, mutational changes in the pathogen that have consequences for infectiousness across different hosts could contribute to evolution towards host specificity (40). Bd can suppress host immune responses and some susceptible species show an over-activation of the immune system that may worsen disease outcomes (41). At the same time, immunization against Bd has been successful in some species, indicating the relationship between adaptive immunity and Bd varies between hosts (42). Amphibian species identity is often the strongest predictor of skin microbial community composition (43,44) and these microbes can be associated with Bd disease outcomes (7,45,46), providing another possible mechanism for facilitating the evolution of host-pathogen specificity. Finally, Bd isolates in the lab can show rapid evolutionary change over short time periods (47,48). Therefore, differences between Bd genotypes in the way they interact with amphibian skin microbes or immune systems, in combination with Bd’s evolutionary lability, could allow for the formation of the host-pathogen genetic correlations that we detected.

In PA, we observed a ubiquitous pattern of association between Bd-GPL1 and bullfrogs in all individual sites. Moreover, not a single bullfrog sample (N=18) in the PA region was infected with Bd-GPL2, or included in the “unassigned” group. The finding that host species was more important than geography in explaining Bd genetic patterns in PA has not been documented in other regions and may indicate that Bd-GPL has been present for a long time in this region and has coevolved with native amphibians, including bullfrogs. We can see additional evidence for this hypothesis at individual sites; at PA Site 4 bullfrogs and the closely related green frog (*Rana clamitans*) were both preferentially infected with Bd-GPL1, while the more distantly related northern leopard frog (*Rana pipiens*) was infected with Bd-GPL2 (49). This pattern follows other studies showing that specialized pathogens are more likely to infect closely related host species (50). Overall, when comparing patterns of Bd genotypes and host species in PA to those documented in other areas of the world, we see evidence for a coevolutionary mechanism shaping Bd-GPL genetic diversity.

An alternative possibility that could produce patterns of host species-Bd genotype correlations is a recent introduction of a novel Bd-GPL lineage into an amphibian community. Existing differences in the ability of the novel Bd-GPL lineage to infect different hosts could maintain the stratification of Bd genotypes along species boundaries. In NV the association between Bd genotypes and host species is not as consistent as in PA, indicating the coexistence of different Bd genotypes in the NV region may be a more recent phenomenon. For example, at NV Site 4 Bd genotypes were strictly partitioned by species, with Bd-GPL1 only present on bullfrogs; however, at NV Site 1 all individuals sampled across species were infected with the same Bd-GPL2 genotype, including bullfrogs. Bullfrogs are not native to Nevada and have been present since at least 1933, likely introduced during fish stocking or from the food trade (51). Therefore, in this region, bullfrog colonization may have provided for a more recent introduction of Bd-GPL1. At NV Site 1 the Bd-GPL2 present may have outcompeted other Bd lineages that could have been present when bullfrogs were first introduced. Supporting this hypothesis, one study found that bullfrogs farmed in Idaho and used for an infection experiment initially harbored cryptic, low intensity infections of Bd-GPL1 that were eventually outcompeted by an inoculation with a high dose of a Bd-GPL2 isolate (52). What host and environmental factors allow for the replacement of one Bd lineage by another is not altogether clear but is a key question going forward to better understand the role of potential disease vectors such the bullfrog.

### What does our data tell us about the origin of Bd-GPL?

The implied coexistence of Bd and amphibians at evolutionary time scales in PA could indicate that the ancestral population of Bd-GPL existed in this part of the world. This possibility has been previously proposed, following reports of higher diversity in Bd samples collected from bullfrog populations (53,54), and evidence that bullfrogs may have facilitated Bd spread from east to west in the US (55). Adding to evidence that North America may be the ancestral source of Bd-GPL, a recent study found as much genetic variation in the Bd-GPL collected from the Sierra Nevada as was present in a large global sample of Bd-GPL genotypes (15). In other regions, such as Panama and Mexico, genetic diversity of Bd-GPL is either spatially structured across the region (Mexico) or genetically homogenous (Panama), the latter indicating a more recent introduction and spread (15,16). Studies using swabs collected from museum specimens have found Bd in Illinois as early as 1888 (56) and in bullfrogs collected in California in 1928 (57). Altogether, a growing body of evidence suggests North America, and perhaps the eastern US (within the native range of the bullfrog) may have been the ancestral population for Bd-GPL. *What can the*

### “unassigned” samples tell us about Bd dynamics?

One key question raised from our data concerns the Bd genotypes that were “unassigned” and lie in between the Bd-GPL1 and Bd-GPL2 genotypes in our PCA. These samples, all of which were from swabs collected in PA, show high levels of heterozygosity across the entire panel of diagnostic alleles. There are two possible explanations for the occurrence of a highly heterozygous, intermediate Bd genotype. The first is that these individuals were coinfected with both Bd-GPL1 and Bd-GPL2. The second is that they were infected with a hybrid lineage resulting from recombination between Bd-GPL1 and Bd-GPL2. Indeed, if we look at the reference Bd isolates in Figure S3 we see that some Bd-GPL2 pure isolates are heterozygous at certain alleles, indicating that high heterozygosity could result from a single infection. Herein, we are unable to confidently say whether these ambiguous samples represent a mixed infection or an infection with a single hybrid lineage.

Our data show not only the coexistence of multiple Bd-GPL lineages at a single site, but also highlights areas where Bd lineage mixing on individuals is rare (in NV, and PA Sites 3 and 4), or common (PA Site 1). The most common frog species caught at PA Site 1 was the green frog (*R. clamitans*), in fact this species accounts for 71% of all unassigned samples in this study. At this point it is unclear whether there is something unique about PA Site 1, or *R. clamitans*, that may allow for such consistent lineage mixing or the potential presence of hybrid lineages. To our knowledge no chytridiomycosis outbreaks have been reported in *R. clamitans*, and individuals from the same site in PA showed no increase in stress hormones when experimentally infected with Bd (58). Probably, this species is a tolerant carrier of Bd, like the closely related bullfrog. This host-pathogen relationship is worth further exploration as the potential for Bd hybridization could lead to novel pathogen phenotypes.

The frequency and circumstances under which recombination occurs in Bd is still unclear. There is, however, plenty of genetic evidence that recombination has occurred between divergent Bd lineages (11,14,59). A likely mechanism for genomic recombination in Bd is parasexual reproduction – where diploid spores fuse to form tetraploid progeny, after which chromosomes are lost as the organism returns back to a diploid state (60). This process has been documented in the human fungal pathogen *Candida albicans*, although the use of this reproductive mode seems to be restricted in nature, possibly to preserve complex gene sets needed to exploit hosts (61). Parasexual reproduction often results in aneuploidy, which is a common phenomenon in Bd (25,62). Furthermore, elevated chromosomal copy number has been linked to increased virulence in Bd (62), and hybrids of Bd have been more deadly than parent lineages when paired with certain host species (63). Indeed, it was originally proposed that Bd-GPL could have originated from recombination between closely related Bd lineages (12), but this hypothesis has been a topic of much debate (64). Most researchers on the subject, however, would agree that precautions should be taken to reduce unintended spread of Bd to limit the possibility of recombination that could produce more deadly lineages.

## Conclusions

Herein we document an association between Bd-GPL genotypes and host species in two regions of the US. We believe the patterns seen in PA may be the result of a long-term coevolutionary relationship between Bd-GPL1 and bullfrogs (and perhaps other closely related species such as the green frog). We observed that the association of Bd-GPL1 with bullfrogs is consistent in NV where bullfrogs have been more recently introduced, and that the patterns in this region are consistent with more recent introductions of novel Bd lineages. We also found evidence of either mixed Bd infections or a potential hybrid lineage that was common at a single site in PA and on green frogs across all PA sites. Given the high genetic diversity and evidence of evolutionary relationships between Bd-GPL lineages and particular host species, we posit that North America may be the ancestral origin of Bd-GPL. Overall, this study presents evidence that Bd-GPL has a long history of coexistence with amphibians in the US, reveals more about the relationship of the bullfrog to Bd, and points to a possible cradle of Bd-GPL genetic diversity in the eastern US.

## Acknowledgements

We thank Frank van Breukelen, Mark Slaughter, and Jonathan Smith for their assistance with funding acquisition and management for the Nevada study. We thank the following people for their assistance with field and laboratory work in Nevada: Rebeca Rivera, Daniel Villanueva, Yesenia Vasquez, David Miller, Joshua Levy, Kevin Guadalupe, Marlai Sai, Ghazal Rezaei, Alexandra Zmuda, Jessica Hill, Shaylene Scarlett, and Alexa Krauss. We thank the following people for assistance with field and laboratory work in Pennsylvania: Laura Brannelly, Karie Altman, Talisin Hammond, Jeffery Bednark, Paradyse Blackwood, Robert Campion, Lauren Chronister, Jordan Coscia, Nina Dunnell, Conor Harrington, Ally Hartman, Marcus Hough, Jennifer Kassimer, Stephanie Kubik, Sadie Parker, Natalie Popielski, Phoebe Reuben, Ayla Ross, Samantha Shablin, Samantha Skerlec, Trina Wantman, Lydia Zimmerman, and Jakub Zegar, Aimee Danley.

## Reproducibility Statement

All data used for this study and R code used for the analyses are available on github: (https://github.com/allie128/Bd_NV_PA).

## Supplementary Figure Legends

**Figure S1:**
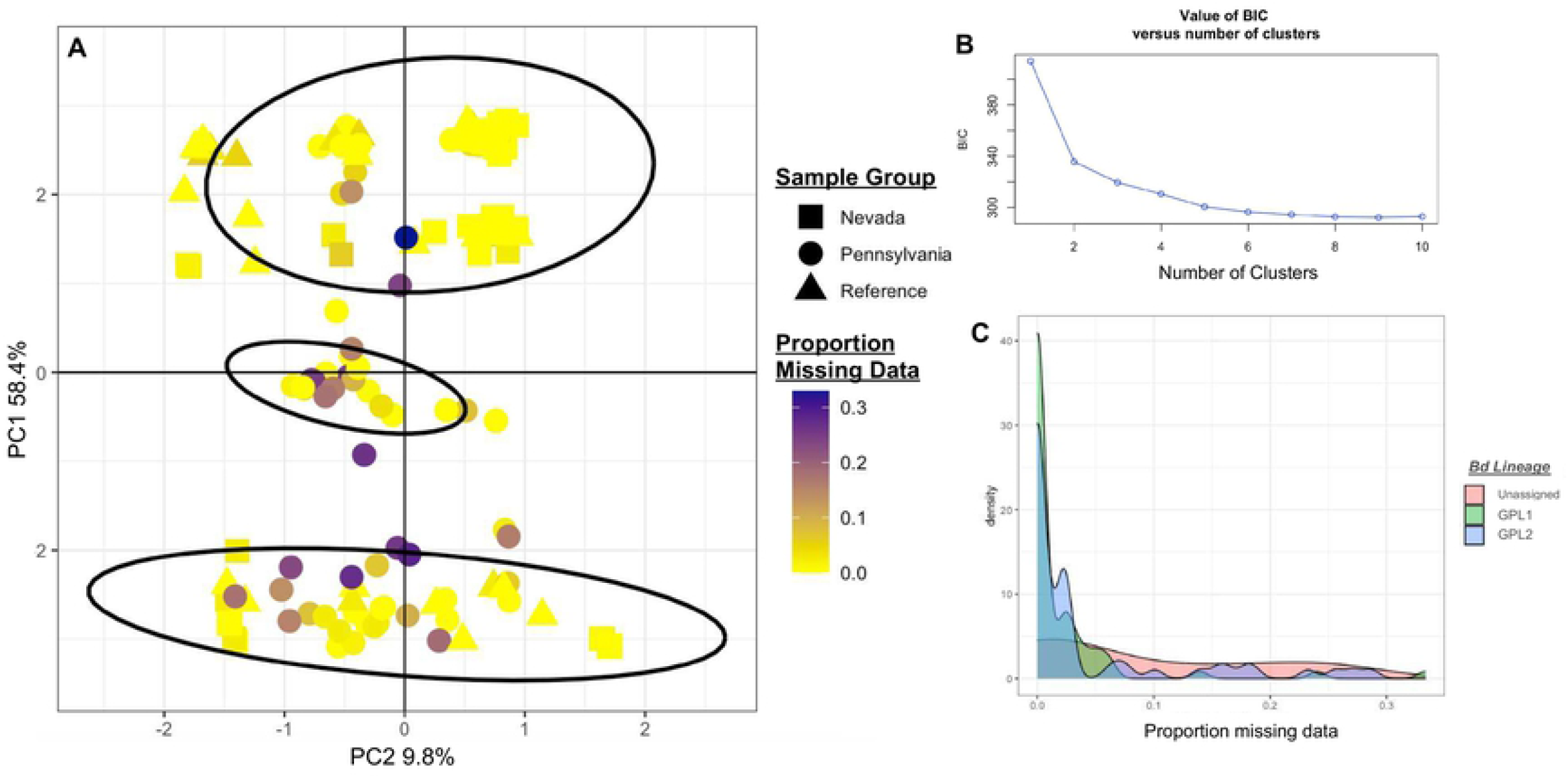
(A) PCA as shown in Figure 1C colored by proportion missing data. (B) BIC chart for DAPC analysis showing K = 2 clusters as the optimal number of clusters (or “elbow” in the chart). (C) Density plot showing that unassigned, Bd-GPL1, and Bd-GPL2 samples all have the same distribution as samples with missing data.

**Figure S2:**
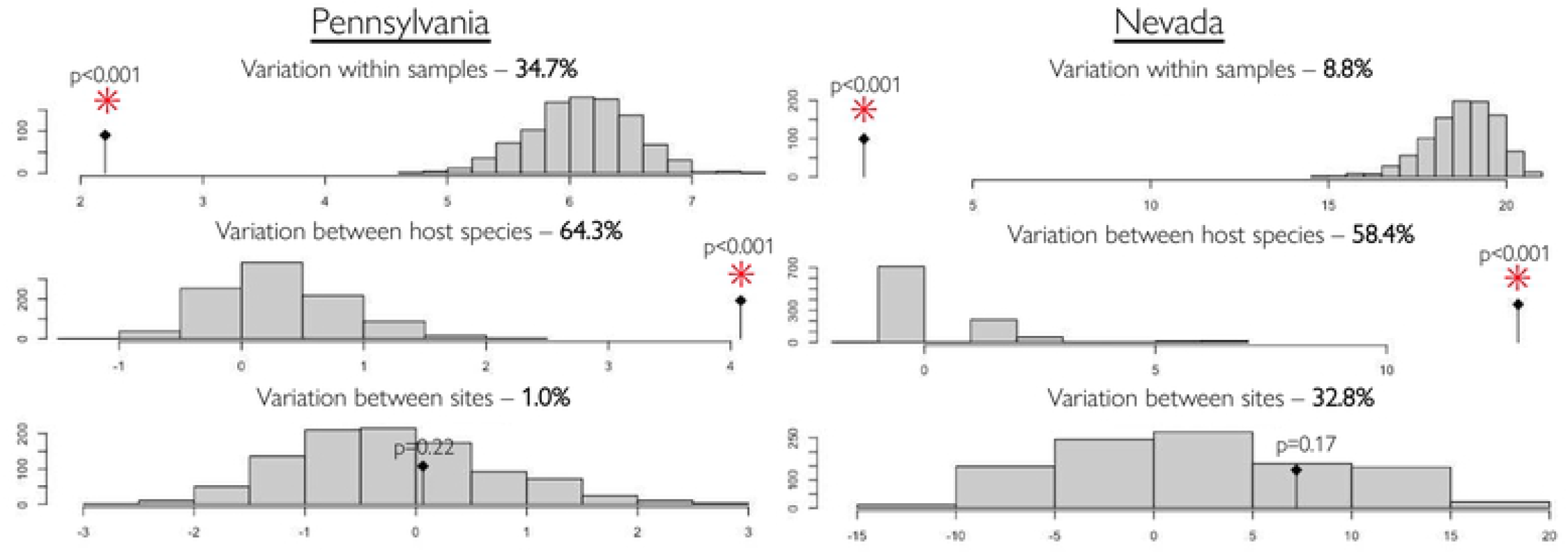
Results from AMOVA testing the relative importance of variation within samples (top), variation between host species (middle), and variation between sites (bottom) in explaining Bd genetic variance for northwestern Pennsylvania (left) and southern Nevada (right). Percentage after each category reflects the relative proportion of variation explained by each category calculated from the components of covariance. Each histogram shows the results from randomly permuting the sample matrices 999 times using the rand.test function in the R package ade4. The black dot indicates the value obtained in this study with a red asterisk indicating statistical significance.

**Figure S3:**
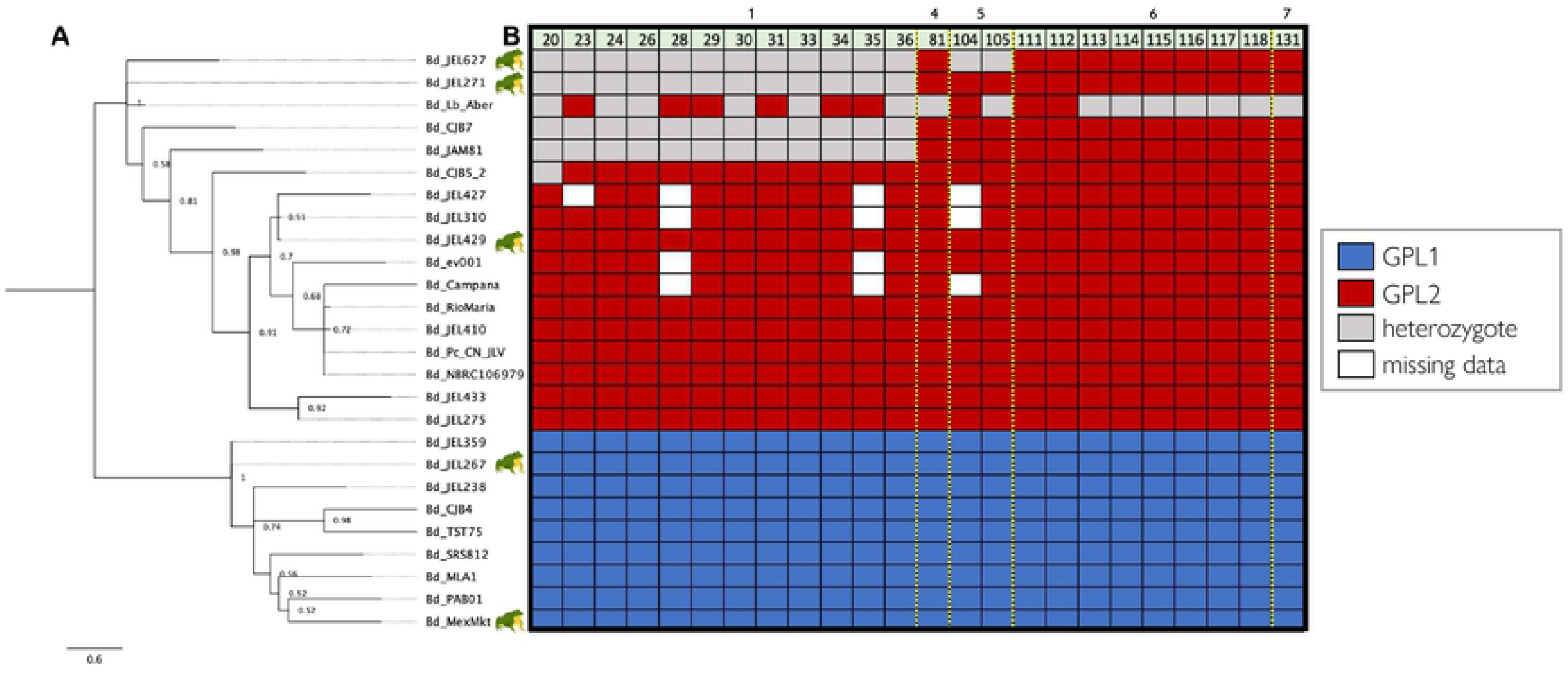
(A) Astral phylogenetic tree and (B) allele plot for 26 pure-isolate Bd-GPL reference samples used to identify the 24 Bd-GPL1 and Bd-GPL2 diagnostic alleles shown in Figures 6 and 7. Tree displayed as in Figures 3 and 4, allele plot displayed as in Figures 6 and 7. A frog icon indicates that isolate was collected from a bullfrog.

## References

1. Kawecki TJ. Red Queen Meets Santa Rosalia: Arms Races and the Evolution of Host Specialization in Organisms with Parasitic Lifestyles. Am Nat [Internet]. 1998 Oct 1;152(4):635–51. Available from: https://doi.org/10.1086/286195

2. Rigaud T, Perrot-Minnot M-J, Brown MJF. Parasite and host assemblages: embracing the reality will improve our knowledge of parasite transmission and virulence. Proc R Soc B Biol Sci [Internet]. 2010 Dec 22;277(1701):3693–702. Available from: https://doi.org/10.1098/rspb.2010.1163

3. Kirchner JW, Roy BA. Evolutionary implications of host–pathogen specificity: fitness consequences of pathogen virulence traits. Evol Ecol Res. 2002;4(1):27–48.

4. Deng X, Gu W, Federman S, du Plessis L, Pybus OG, Faria NR, et al. Genomic surveillance reveals multiple introductions of SARS-CoV-2 into Northern California. Science (80-) [Internet]. 2020 Jul 31;369(6503):582LP–587. Available from: http://science.sciencemag.org/content/369/6503/582.abstract

5. Longcore JE, Pessier AP, Nichols DK. Batrachochytrium Dendrobatidis gen. et sp. nov., a Chytrid Pathogenic to Amphibians. Mycologia [Internet]. 1999;91(2):219–27. Available from: http://www.jstor.org/stable/3761366

6. Lips KR, Diffendorfer J, Mendelson III JR, Sears MW. Riding the Wave: Reconciling the Roles of Disease and Climate Change in Amphibian Declines. PLoS Biol [Internet]. 2008 Mar 25;6(3):e72. Available from: http://dx.doi.org/10.1371%2Fjournal.pbio.0060072

7. Woodhams DC, Voyles J, Lips KR, Carey C, Rollins-Smith LA. PREDICTED DISEASE SUSCEPTIBILITY IN A PANAMANIAN AMPHIBIAN ASSEMBLAGE BASED ON SKIN PEPTIDE DEFENSES. J Wildl Dis [Internet]. 2006 Apr 1;42(2):207–18. Available from: https://doi.org/10.7589/0090-3558-42.2.207

8. Ron SR, Merino A. Amphibian declines in Ecuador: overview and first report of chytridiomycosis from South America. Froglog. 2000;42:2–3.

9. Briggs CJ, Vredenburg VT, Knapp RA, Rachowicz LJ. Investigating the Population-Level effects of Chytridiomycosis: An Emerging Infectious Disease of Amphibians. Ecology [Internet]. 2005 Dec 1;86(12):3149–59. Available from: http://dx.doi.org/10.1890/04-1428

10. Berger L, Speare R, Daszak P, Green DE, Cunningham AA, Goggin CL, et al. Chytridiomycosis causes amphibian mortality associated with population declines in the rain forests of Australia and Central America. Proc Natl Acad Sci [Internet]. 1998 Jul 21;95(15):9031–6. Available from: http://www.pnas.org/content/95/15/9031.abstract

11. O’Hanlon SJ, Rieux A, Farrer RA, Rosa GM, Waldman B, Bataille A, et al. Recent Asian origin of chytrid fungi causing global amphibian declines. Science (80-) [Internet]. 2018 May 11;360(6389):621LP–627. Available from: http://science.sciencemag.org/content/360/6389/621.abstract

12. Farrer RA, Weinert LA, Bielby J, Garner TWJ, Balloux F, Clare F, et al. Multiple emergences of genetically diverse amphibian-infecting chytrids include a globalized hypervirulent recombinant lineage. Proc Natl Acad Sci [Internet]. 2011 Nov 15;108(46):18732–6. Available from: http://www.pnas.org/content/108/46/18732.abstract

13. Byrne AQ, Vredenburg VT, Martel A, Pasmans F, Bell RC, Blackburn DC, et al. Cryptic diversity of a widespread global pathogen reveals expanded threats to amphibian conservation. Proc Natl Acad Sci [Internet]. 2019 Oct 8;116(41):20382LP–20387. Available from: http://www.pnas.org/content/116/41/20382.abstract

14. Schloegel LM, Toledo LF, Longcore JE, Greenspan SE, Vieira CA, Lee M, et al. Novel, panzootic and hybrid genotypes of amphibian chytridiomycosis associated with the bullfrog trade. Mol Ecol [Internet]. 2012;21(21):5162–77. Available from: http://www.ncbi.nlm.nih.gov/pubmed/22857789

15. Rothstein AP, Byrne AQ, Knapp RA, Briggs CJ, Voyles J, Richards-Zawacki CL, et al. Divergent regional evolutionary histories of a devastating global amphibian pathogen. Proc R Soc B. 2021;288(1953):20210782.

16. Basanta MD, Byrne AQ, Rosenblum EB, Piovia-Scott J, Parra-Olea G. Early presence of Batrachochytrium dendrobatidis in Mexico with a contemporary dominance of the global panzootic lineage. Mol Ecol. 2021;30(2):424–37.

17. Phillott AD, Speare R, Hines HB, Skerratt LF, Meyer E, McDonald KR, et al. Minimising exposure of amphibians to pathogens during field studies. Dis Aquat Organ. 2010;92(2–3):175–85.

18. Jaeger JR, Waddle AW, Rivera R, Harrison DT, Ellison S, Forrest MJ, et al. Batrachochytrium dendrobatidis and the Decline and Survival of the Relict Leopard Frog. Ecohealth. 2017;14(2):285–95.

19. Boyle DG, Boyle DB, Olsen V, Morgan JAT, Hyatt AD. Rapid quantitative detection of chytridiomycosis (Batrachochytrium dendrobatidis) in amphibian samples using real-time Taqman PCR assay. 2004;60:141–8. Available from: https://dx.doi.org/10.3354/dao060141

20. Waddle AW, Levy JE, Rivera R, van Breukelen F, Nash M, Jaeger JR. Population-level resistance to chytridiomycosis is life-stage dependent in an imperiled anuran. Ecohealth. 2019;16(4):701–11.

21. Waddle AW, Sai M, Levy JE, Rezaei G, van Breukelen F, Jaeger JR. Systematic approach to isolating Batrachochytrium dendrobatidis. Dis Aquat Organ. 2018;127(3):243–7.

22. Zolan ME, Pukkila PJ. Inheritance of DNA methylation in Coprinus cinereus. Mol Cell Biol [Internet]. 1986 Jan 1;6(1):195–200. Available from: http://mcb.asm.org/content/6/1/195.abstract

23. Byrne AQ, Rothstein AP, Poorten TJ, Erens J, Settles ML, Rosenblum EB. Unlocking the story in the swab: A new genotyping assay for the amphibian chytrid fungus Batrachochytrium dendrobatidis. Mol Ecol Resour [Internet]. 2017 May 3 [cited 2017 Jul 21]; Available from: http://doi.wiley.com/10.1111/1755-0998.12675

24. Martel A, Spitzen-van der Sluijs A, Blooi M, Bert W, Ducatelle R, Fisher MC, et al. Batrachochytrium salamandrivorans sp. nov. causes lethal chytridiomycosis in amphibians. Proc Natl Acad Sci [Internet]. 2013 Sep 17;110(38):15325–9. Available from: http://www.pnas.org/content/110/38/15325.abstract

25. Rosenblum EB, James TY, Zamudio KR, Poorten TJ, Ilut D, Rodriguez D, et al. Complex history of the amphibian-killing chytrid fungus revealed with genome resequencing data. Proc Natl Acad Sci USA [Internet]. 2013;110(23):9385–90. Available from: http://www.pubmedcentral.nih.gov/articlerender.fcgi?artid=3677446&tool=pmcentrez&rendertype=abstract

26. Edgar RC. MUSCLE: multiple sequence alignment with high accuracy and high throughput. Nucleic Acids Res [Internet]. 2004 Jan 19 [cited 2016 Sep 2];32(5):1792–7. Available from: http://nar.oxfordjournals.org/cgi/content/long/32/5/1792

27. Kearse M, Moir R, Wilson A, Stones-Havas S, Cheung M, Sturrock S, et al. Geneious Basic: An integrated and extendable desktop software platform for the organization and analysis of sequence data. Bioinforma [Internet]. 2012 Jun 15;28(12):1647–9. Available from: http://bioinformatics.oxfordjournals.org/content/28/12/1647.abstract

28. Stamatakis A. RAxML version 8: a tool for phylogenetic analysis and post-analysis of large phylogenies. Bioinformatics [Internet]. 2014 May 1;30(9):1312–3. Available from: http://dx.doi.org/10.1093/bioinformatics/btu033

29. Zhang C, Rabiee M, Sayyari E, Mirarab S. ASTRAL-III: polynomial time species tree reconstruction from partially resolved gene trees. BMC Bioinformatics [Internet]. 2018;19(6):153. Available from: https://doi.org/10.1186/s12859-018-2129-y

30. Li H, Durbin R. Fast and accurate short read alignment with Burrows–Wheeler transform. Bioinformatics [Internet]. 2009 May 18;25(14):1754–60. Available from: https://doi.org/10.1093/bioinformatics/btp324

31. Garrison E, Marth G. Haplotype-based variant detection from short-read sequencing. arXiv Prepr arXiv12073907. 2012;

32. Danecek P, Auton A, Abecasis G, Albers CA, Banks E, DePristo MA, et al. The variant call format and VCFtools. Bioinformatics [Internet]. 2011 Aug 1;27(15):2156–8. Available from: https://doi.org/10.1093/bioinformatics/btr330

33. Knaus BJ, Grünwald NJ. vcfr: a package to manipulate and visualize variant call format data in R. Mol Ecol Resour [Internet]. 2017 Jan 1;17(1):44–53. Available from: https://doi.org/10.1111/1755-0998.12549

34. Jombart T, Devillard S, Balloux F. Discriminant analysis of principal components: a new method for the analysis of genetically structured populations. BMC Genet [Internet]. 2010;11(1):94. Available from: https://doi.org/10.1186/1471-2156-11-94

35. Jombart T. adegenet: a R package for the multivariate analysis of genetic markers. Bioinformatics. 2008;24(11):1403–5.

36. Excoffier L, Smouse PE, Quattro JM. Analysis of molecular variance inferred from metric distances among DNA haplotypes: application to human mitochondrial DNA restriction data. Genetics. 1992;131(2):479–91.

37. Kamvar ZN, Tabima JF, Grünwald NJ. Poppr: an R package for genetic analysis of populations with clonal, partially clonal, and/or sexual reproduction. PeerJ. 2014;2:e281.

38. Voyles J, Woodhams DC, Saenz V, Byrne AQ, Perez R, Rios-Sotelo G, et al. Shifts in disease dynamics in a tropical amphibian assemblage are not due to pathogen attenuation. Science (80-) [Internet]. 2018 Mar 30;359(6383):1517LP–1519. Available from: http://science.sciencemag.org/content/359/6383/1517.abstract

39. James TY, Toledo LF, Rödder D, da Silva Leite D, Belasen AM, Betancourt-Román CM, et al. Disentangling host, pathogen, and environmental determinants of a recently emerged wildlife disease: lessons from the first 15 years of amphibian chytridiomycosis research. Ecol Evol [Internet]. 2015;n/a-n/a. Available from: http://doi.wiley.com/10.1002/ece3.1672

40. Woolhouse MEJ, Taylor LH, Haydon DT. Population Biology of Multihost Pathogens. Science (80-) [Internet]. 2001 May 11;292(5519):1109–12. Available from: https://doi.org/10.1126/science.1059026

41. Ellison AR, Savage AE, DiRenzo G V, Langhammer P, Lips KR, Zamudio KR. Fighting a Losing Battle: Vigorous Immune Response Countered by Pathogen Suppression of Host Defenses in the Chytridiomycosis-Susceptible Frog Atelopus zeteki. G3 (Bethesda) [Internet]. 2014;4(July):1275–89. Available from: http://www.ncbi.nlm.nih.gov/pubmed/24841130

42. Waddle AW, Rivera R, Rice H, Keenan EC, Rezaei G, Levy JE, et al. Amphibian resistance to chytridiomycosis increases following low-virulence chytrid fungal infection or drug-mediated clearance. J Appl Ecol. 2021;1–12.

43. Kueneman JG, Parfrey LW, Woodhams DC, Archer HM, Knight R, McKenzie VJ. The amphibian skin-associated microbiome across species, space and life history stages. Mol Ecol [Internet]. 2014 Mar 1;23(6):1238–50. Available from: https://doi.org/10.1111/mec.12510

44. McKenzie VJ, Bowers RM, Fierer N, Knight R, Lauber CL. Co-habiting amphibian species harbor unique skin bacterial communities in wild populations. ISME J [Internet]. 2012;6(3):588–96. Available from: https://doi.org/10.1038/ismej.2011.129

45. Lam BA, Walke JB, Vredenburg VT, Harris RN. Proportion of individuals with anti-Batrachochytrium dendrobatidis skin bacteria is associated with population persistence in the frog Rana muscosa. Biol Conserv [Internet]. 2010;143(2):529–31. Available from: http://www.sciencedirect.com/science/article/pii/S0006320709004807

46. Piovia-Scott J, Rejmanek D, Woodhams DC, Worth SJ, Kenny H, McKenzie V, et al. Greater Species Richness of Bacterial Skin Symbionts Better Suppresses the Amphibian Fungal Pathogen Batrachochytrium Dendrobatidis. Microb Ecol [Internet]. 2017;74(1):217–26. Available from: https://doi.org/10.1007/s00248-016-0916-4

47. Langhammer PF, Lips KR, Burrowes PA, Tunstall T, Palmer CM, Collins JP. A Fungal Pathogen of Amphibians, Batrachochytrium dendrobatidis, Attenuates in Pathogenicity with In Vitro Passages. PLoS One [Internet]. 2013 Oct 10;8(10):e77630. Available from: http://dx.doi.org/10.1371%2Fjournal.pone.0077630

48. Refsnider JM, Poorten TJ, Langhammer PF, Burrowes PA, Rosenblum EB. Genomic Correlates of Virulence Attenuation in the Deadly Amphibian Chytrid Fungus, Batrachochytrium dendrobatidis. 2015;5(November):2291–8. Available from: https://doi.org/10.1534/g3.115.021808

49. Yuan Z-Y, Zhou W-W, Chen X, Poyarkov Nikolay A. J, Chen H-M, Jang-Liaw N-H, et al. Spatiotemporal Diversification of the True Frogs (Genus Rana): A Historical Framework for a Widely Studied Group of Model Organisms. Syst Biol [Internet]. 2016 Sep 1;65(5):824–42. Available from: http://dx.doi.org/10.1093/sysbio/syw055

50. Davies TJ, Pedersen AB. Phylogeny and geography predict pathogen community similarity in wild primates and humans. Proc R Soc B Biol Sci [Internet]. 2008 Jul 22;275(1643):1695–701. Available from: https://doi.org/10.1098/rspb.2008.0284

51. Linsdale JM. Amphibians and Reptiles in Nevada. Proc Am Acad Arts Sci [Internet]. 1940 Oct 22;73(8):197–257. Available from: http://www.jstor.org/stable/25130182

52. Byrne AQ, Poorten TJ, Voyles J, Willis CKR, Rosenblum EB. Opening the file drawer: Unexpected insights from a chytrid infection experiment. PLoS One. 2018;13(5):e0196851.

53. James TY, Litvintseva AP, Vilgalys R, Morgan JAT, Taylor JW, Fisher MC, et al. Rapid Global Expansion of the Fungal Disease Chytridiomycosis into Declining and Healthy Amphibian Populations. PLOS Pathog [Internet]. 2009 May 29;5(5):e1000458. Available from: https://doi.org/10.1371/journal.ppat.1000458

54. Goka K, Yokoyama J, Une Y, Kuroki T, Suzuki K, Nakahara M, et al. Amphibian chytridiomycosis in Japan: distribution, haplotypes and possible route of entry into Japan. Mol Ecol [Internet]. 2009 Dec [cited 2016 Aug 22];18(23):4757–74. Available from: http://www.ncbi.nlm.nih.gov/pubmed/19840263

55. Yap TA, Koo MS, Ambrose RF, Vredenburg VT. Introduced bullfrog facilitates pathogen invasion in the western United States. PLoS One [Internet]. 2018 Apr 16;13(4):e0188384. Available from: https://doi.org/10.1371/journal.pone.0188384

56. Talley BL, Muletz CR, Vredenburg VT, Fleischer RC, Lips KR. A century of Batrachochytrium dendrobatidis in Illinois amphibians (1888–1989). Biol Conserv [Internet]. 2015;182(October):254–61. Available from: http://linkinghub.elsevier.com/retrieve/pii/S0006320714004807

57. Huss M, Huntley L, Vredenburg V, Johns J, Green S. Prevalence of Batrachochytrium dendrobatidis in 120 archived specimens of Lithobates catesbeianus (American bullfrog) collected in California, 1924-2007. Ecohealth. 2013;10(4):339–43.

58. Hammond TT, Blackwood PE, Shablin SA, Richards-Zawacki CL. Relationships between glucocorticoids and infection with Batrachochytrium dendrobatidis in three amphibian species. Gen Comp Endocrinol [Internet]. 2020;285:113269. Available from: https://www.sciencedirect.com/science/article/pii/S0016648019302953

59. Jenkinson TS, Betancourt Román CM, Lambertini C, Valencia-Aguilar A, Rodriguez D, Nunes-de-Almeida CHL, et al. Amphibian-killing chytrid in Brazil comprises both locally endemic and globally expanding populations. Mol Ecol [Internet]. 2016 Jul [cited 2016 Aug 26];25(13):2978–96. Available from: http://www.ncbi.nlm.nih.gov/pubmed/26939017

60. Samarasinghe H, You M, Jenkinson TS, Xu J, James TY. Hybridization Facilitates Adaptive Evolution in Two Major Fungal Pathogens. Vol. 11, Genes. 2020.

61. Nielsen K, Heitman JBT-A in G. Sex and Virulence of Human Pathogenic Fungi. In: Fungal Genomics [Internet]. Academic Press. 2007. p. 143–73. Available from: https://www.sciencedirect.com/science/article/pii/S006526600657004X

62. Farrer RA, Henk DA, Garner TWJ, Balloux F, Woodhams DC, Fisher MC. Chromosomal copy number variation, selection and uneven rates of recombination reveal cryptic genome diversity linked to pathogenicity. PLoS Genet. 2013;9(8):e1003703.

63. Greenspan SE, Lambertini C, Carvalho T, James TY, Toledo LF, Haddad CFB, et al. Hybrids of amphibian chytrid show high virulence in native hosts. Sci Rep [Internet]. 2018;8(1):9600. Available from: https://doi.org/10.1038/s41598-018-27828-w

64. Fisher MC, Garner TWJ. Chytrid fungi and global amphibian declines. Nat Rev Microbiol [Internet]. 2020;18(6):332–43. Available from: https://doi.org/10.1038/s41579-020-0335-x

